# Monocyte-derived transcriptome signature indicates antibody-dependent cellular phagocytosis as the primary mechanism of vaccine-induced protection against HIV-1

**DOI:** 10.1101/2021.05.12.443737

**Authors:** Shida Shangguan, Philip K. Ehrenberg, Aviva Geretz, Lauren Yum, Gautam Kundu, Kelly May, Slim Fourati, Krystelle Nganou-Makamdop, LaTonya D. Williams, Sheetal Sawant, Eric Lewitus, Punnee Pitisuttithum, Sorachai Nitayaphan, Suwat Chariyalertsak, Supachai Rerks-Ngarm, Morgane Rolland, Daniel Douek, Peter Gilbert, Georgia D. Tomaras, Nelson Michael, Sandhya Vasan, Rasmi Thomas

## Abstract

A gene signature previously correlated with mosaic adenovirus 26 vaccine protection in simian immunodeficiency virus (SIV) and SHIV challenge models in non-human primates (NHP). In this report we investigated presence of this signature as a correlate of reduced risk in human clinical trials and potential mechanism for protection. The absence of this gene signature in the DNA/rAd5 human vaccine trial which did not show efficacy, strengthens our hypothesis that this signature is only enriched in studies that demonstrated protection. This gene signature was enriched in the partially effective RV144 human trial that administered the ALVAC/protein vaccine, and we find that the signature associates with both decreased risk of HIV-1 acquisition and increased vaccine efficacy. Total RNA-seq in a clinical trial that used the same vaccine regimen as the RV144 HIV vaccine implicated antibody-dependent cellular phagocytosis (ADCP) as a potential mechanism of vaccine protection. CITE-seq profiling of 53 surface markers and transcriptomes of 53,777 single cells from the same trial, showed that genes in this signature were primarily expressed in cells belonging to the myeloid lineage including monocytes, which are major effector cells for ADCP. The consistent association of this transcriptome signature with vaccine efficacy represents a tool both to identify potential mechanisms, as with ADCP here, and to screen novel approaches to accelerate development of new vaccine candidates.

## Introduction

HIV vaccines are being tested in ongoing HIV human efficacy trials including the HVTN 705 and 706 studies (Hsu & O’Connell, 2017; Mega, 2019). Vaccines from these trials were previously tested in non-human primate (NHP) and showed partial protection from infection (Barouch et al., 2015; Barouch et al., 2018). These vaccines are based on the adenovirus serotype 26 (Ad26)-based regimen and have been tested in human immunogenicity trials (Baden et al., 2020; Barouch et al., 2018). To date, the pivotal RV144 phase 3 human efficacy trial conducted in Thailand is the only vaccine to show any protection against HIV (Rerks-Ngarm et al., 2009). The vaccine used a canary-pox ALVAC-based vector with a bivalent gp120 protein boost. Although neither of the vaccine regimens using different viral vectors were fully efficacious, there is some consensus that current preventive and treatment methods along with a moderately effective vaccine could potentially reduce the HIV pandemic (Anderson, Swinton, & Garnett, 1995; Andersson et al., 2007; Fauci, 2017; Medlock et al., 2017). While a number of correlates of vaccine protection have been described for these studies, protection mediated by the humoral immune system including HIV-1 specific IgG antibody titers, antibody Fc polyfunctionality, antibody interactions with HLA class II gene products and antibody effector functions have been key features of these partially effective vaccines (Barouch et al., 2015; Barouch et al., 2018; Haynes et al., 2012; Prentice et al., 2015).

Recently, we showed that a vaccine-induced gene signature identified in B cells by an unbiased transcriptome-wide RNA-seq approach associated with decreased risk against SIV/SHIV infection in NHP studies evaluating the Ad26 vaccine (Ehrenberg et al., 2019). This geneset was also enriched in NHP and the human RV144 trial that employed a vaccine containing the ALVAC viral vector (Ehrenberg et al., 2019). This gene signature is not merely a general response elicited by vaccination as it was not enriched in the Ad26-MVA arm of the SHIV challenge in NHP that showed some protection previously (Barouch et al., 2018). This gene signature was initially defined when comparing differentially expressed genes between B cells and monocytes from vaccinated individuals in an Influenza immunogenicity trial (Nakaya et al., 2011). The geneset that was submitted to the molecular signature database (MSigdb) comprised the top 200 genes that were upregulated in monocytes compared to B cells. In our previous study, specific genes in this geneset were upregulated in the uninfected compared to infected group in multiple SIV/HIV trials (Ehrenberg et al., 2019). Genes that were previously correlated with immunogenicity in human vaccine trials of influenza and yellow fever, including TNFSF13 (APRIL), were also enriched in uninfected rhesus monkeys in the NHP studies (Li et al., 2014; Nakaya et al., 2011). Although we first identified this geneset in sorted B cells (Ehrenberg et al., 2019), we were able to identify the same signature associating with reduced infection in published microarray datasets from both bulk unstimulated and *in vitro* antigen-stimulated PBMCs from three independent preclinical and clinical studies (Fourati et al., 2019; Vaccari et al., 2018; Vaccari et al., 2016). To determine if specific immune responses might be driving protection in conjunction with the gene signature, we examined whether this geneset associated with these responses measured in the NHP studies. We observed that this gene signature was also enriched in animals with increased magnitude of ADCP (Ehrenberg et al., 2019). We propose that this gene signature is a correlate of reduced risk of infection in efficacy studies and further investigation of the enriched genes in the geneset could potentially help uncover the mechanism of vaccine protection. Here we investigate this gene signature further as a proxy of vaccine-induced protection in human clinical trials, to identify the cellular origin, as well as to investigate potential mechanisms for the decreased risk of infection.

## Results

### Gene signature is absent in a human HIV vaccine trial that did not show efficacy

Since the gene signature associated with vaccine protection in multiple studies using different regimens, we wanted to further confirm that this signature was truly associated with protection by looking for its presence (or absence) in a human vaccine trial that failed to show efficacy. We screened for this gene signature in whole-transcriptome data from participants vaccinated with the DNA/rAd5 HIV-1 preventive vaccine in the HVTN 505 human efficacy trial. Immunizations in this trial were halted prior to reaching the clinical endpoint due to lack of efficacy (Hammer et al., 2013). When comparing infection status, enrichment of this gene signature, as defined by the normalized enrichment score (NES), was not significant in transcriptome data from sorted B cells or monocytes one month after the final immunization in the vaccinated arm of the study (NES=−1.18, P=0.09 and NES=1.12, P=0.18 respectively). This finding further supports our hypothesis that this gene signature is associated with protection as summarized in Table 1.

**Table 1.** Gene signature associates with vaccine protection in multiple trials.

| Study | Vaccine Regimen | Species | Partial protection | N | Method | Protective signature |
| --- | --- | --- | --- | --- | --- | --- |
| 09-11 | Ad26/gp140 | NHP | Y | 10 | RNA-seq | Y |
| 13-19 | Ad26/gp140 | NHP | Y | 11 | RNA-seq | Y |
| 13-19 | A26/Ad26 +gp140 | NHP | Y | 12 | RNA-seq | Y |
| 13-19 | Ad26/MVA+gp140 | NHP | Y | 9 | RNA-seq | N |
|  | ALVAC-SIV/gp120 | NHP | Y | 27 | Microarray | Y |
|  | DNA-SIV/ALVAC+gp120 | NHP | Y | 12 | Microarray | Y |
| RV144 | ALVAC/gp120 | Human | Y | 170 | Microarray | Y |
| HVTN 505 | DNA/rAd5 | Human | N | 42 | RNA-seq | N |

### GES is the strongest correlate of protection in RV144

In previous analyses of NHP preclinical studies, we utilized a composite gene expression score (GES) consisting of an average of standardized expression of the specific number of enriched genes in one study, to predict infection status in an independent study using the overlapping expressed genes from the first study (Ehrenberg et al., 2019). While this method is successful in evaluating gene signatures in studies using similar vaccine strategies, we wanted to explore this approach across different studies using diverse platforms. For each independent study, we computed a GES derived from genes within the geneset that were enriched in uninfected donors by averaging standardized expression and showed that it associates with decreased HIV-1 infection (Fig. 1). The magnitude of the GES and total number of enriched genes present in the gene signature is specific to each study and is higher in the uninfected animals in the two NHP preclinical trials evaluating the mosaic Ad26 vaccine (09-11 and 13-19), including the different arms of the 13-19 study (13-19a-b) (Fig. 1A-C). Further, in the RV144 trial, the GES of 63 enriched genes in the gene signature also was higher in the vaccinated individuals that remained uninfected (Fig. 1D). The number of enriched genes in each study might vary due to global differences in the vaccine strategies, but we consistently observed that higher GES associated with protection from HIV acquisition. We took advantage of the composite GES measurement to compare it with the other known primary correlates of HIV-1 infection risk in the human RV144 trial. IgG antibodies binding to the variable regions 1-2 (V1V2) of the HIV-1 Envelope (Env) have been shown to correlate with decreased risk of infection, while IgA binding to Env associated with increased risk of infection (Haynes et al., 2012). We show that the association of the GES in RV144 is a stronger correlate of reduced risk of infection than the previously described V1V2-specific IgG antibodies (Fig. 2A). Cumulative incidence curves of HIV-1 infection showed decreased rates of infection among vaccine recipients with high GES (Fig. 2B) Estimated vaccine efficacy was higher among vaccine recipients with higher GES (Fig. 2C). The distribution of area under the receiver operating characteristic curve (AUC) and accuracy suggested that GES was also able to predict HIV-1 infection (Fig. 2D). The effect of GES was also tested in RV144 vaccine and placebo participants who became infected during the trial (Rolland et al., 2012). If the GES was associated with vaccine efficacy, we would expect that vaccinees with a high GES did not get infected, hence vaccinees who became infected should have lower GES than placebo participants (who reflect the entire distribution of GES). This was observed across 43 breakthrough participants, with a significant difference among participants infected with single HIV-1 founder variants (N=29) (Supplementary Fig. 1). These findings strengthen the hypothesis that the GES is associated with vaccine efficacy.

**Fig. 1.**
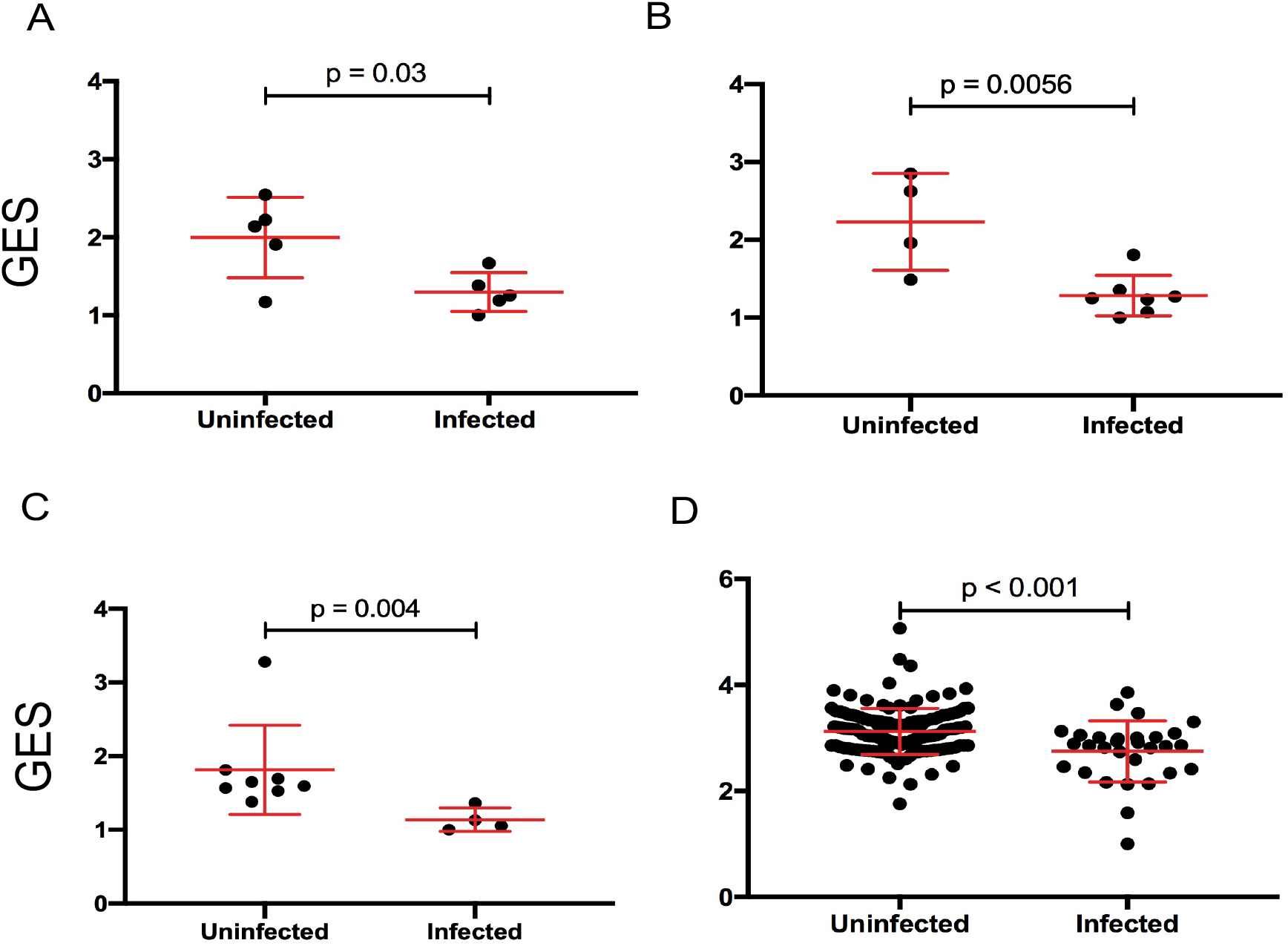
Composite gene expression scores (GES) are higher in the uninfected compared to infected groups. GES computed from enriched genes in the geneset is higher in the uninfected compared to infected vaccinated NHP and humans. (A) Ad26/gp140 (09-11 NHP SIV challenge study, 58 enriched genes, N = 10), (B) Ad26/gp140 (13-19 NHP SHIV challenge study, 58 enriched genes, N = 11), (C) Ad26/Ad26+gp140 (13-19 NHP SHIV challenge study, 68 enriched genes, N = 12), and (D) ALVAC/gp 120 (RV144 human efficacy trial, 63 enriched genes, N = 170). The statistical significance was calculated by either Mann-Whitney or unpaired t-test.

**Fig. 2.**
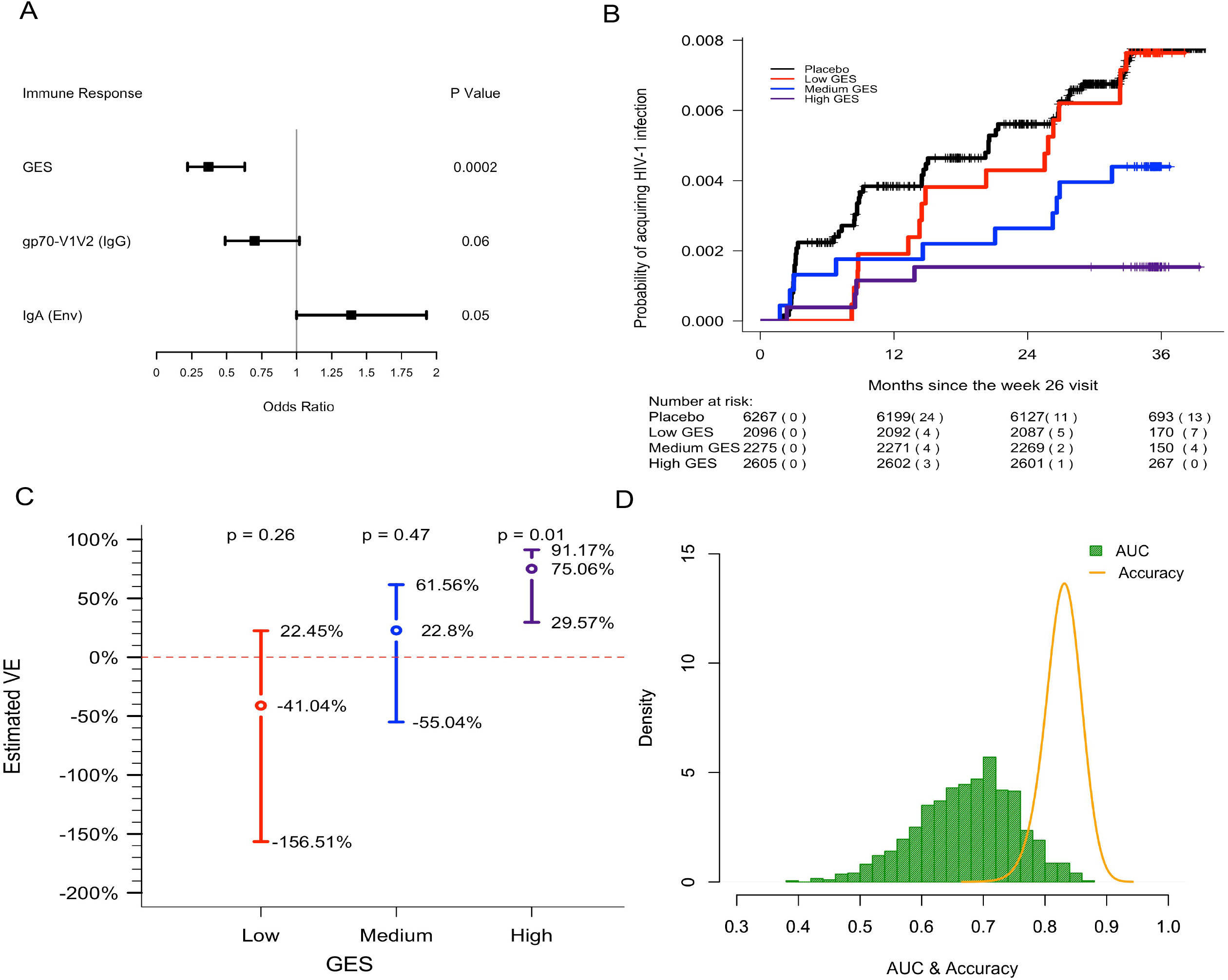
GES is a stronger correlate of reduced risk of infection in RV144. A GES of the 63 enriched genes in the RV144 study was examined as a continuous variable (N = 170). (A) GES is associated with lower odds of HIV acquisition compared to the other two primary correlates of risk. Variables were measured at week 26, two weeks post last vaccination. For each variable, the odds ratio is reported per 1-SD increase. Transcriptome data was available only in a subset of the 246 donors. (B) Probability of acquiring HIV-1 is lower in individuals with higher GES. (C) Vaccine efficacy is increased significantly in individuals with high GES. (D) Distribution of AUC and accuracy plotted after repeating the process 1000 times showed that GES could predict HIV-1 infection with AUC of 0.67 ± 0.08 and with accuracy of 0.83 ± 0.02.

### Gene signature associates with an antibody effector function in a human vaccine trial

Immune responses correlating with this signature can provide additional insights into mechanisms that could be harnessed to improve vaccine design. We previously showed in the NHP studies that the protective gene signature that was enriched in uninfected monkeys after Ad26/gp140 vaccination also associated with higher magnitude of ADCP (Ehrenberg et al., 2019). In the RV144 human trial a number of immunological parameters were previously measured as part of the immune-correlates analysis, but not ADCP. The RV306 immunogenicity trial that employed a similar prime boost RV144 vaccine regimen with additional late boosts provided us with a unique opportunity to test if the gene signature was associated with ADCP (Pitisuttithum et al., 2020). We generated transcriptome-wide gene expression data from peripheral blood two weeks after the RV144 vaccine regimen (prior to the additional boosts) and assessed for enrichment of the gene signature with the magnitude of ADCP measured at the same timepoint in 24 participants. The gene signature with 118 enriched genes was significantly associated with higher magnitude of ADCP (NES=3.0, P<0.001)(Fig. 3A). Using the same geneset, 93 genes were found to be enriched in a subset of overlapping participants (N=21), where samples were collected 3 days after the RV144 immunizations (NES=2.5, P<0.001)(Fig. 3B). A GES from the list of enriched genes associating with ADCP from both timepoints was computed in the RV144 dataset. ADCP GES from both timepoints correlated strongly with the protective RV144 GES (rho=0.74, P= 2.2e-16, rho=0.75, P= 2.2e-16)(Fig. 3C).

**Fig. 3.**
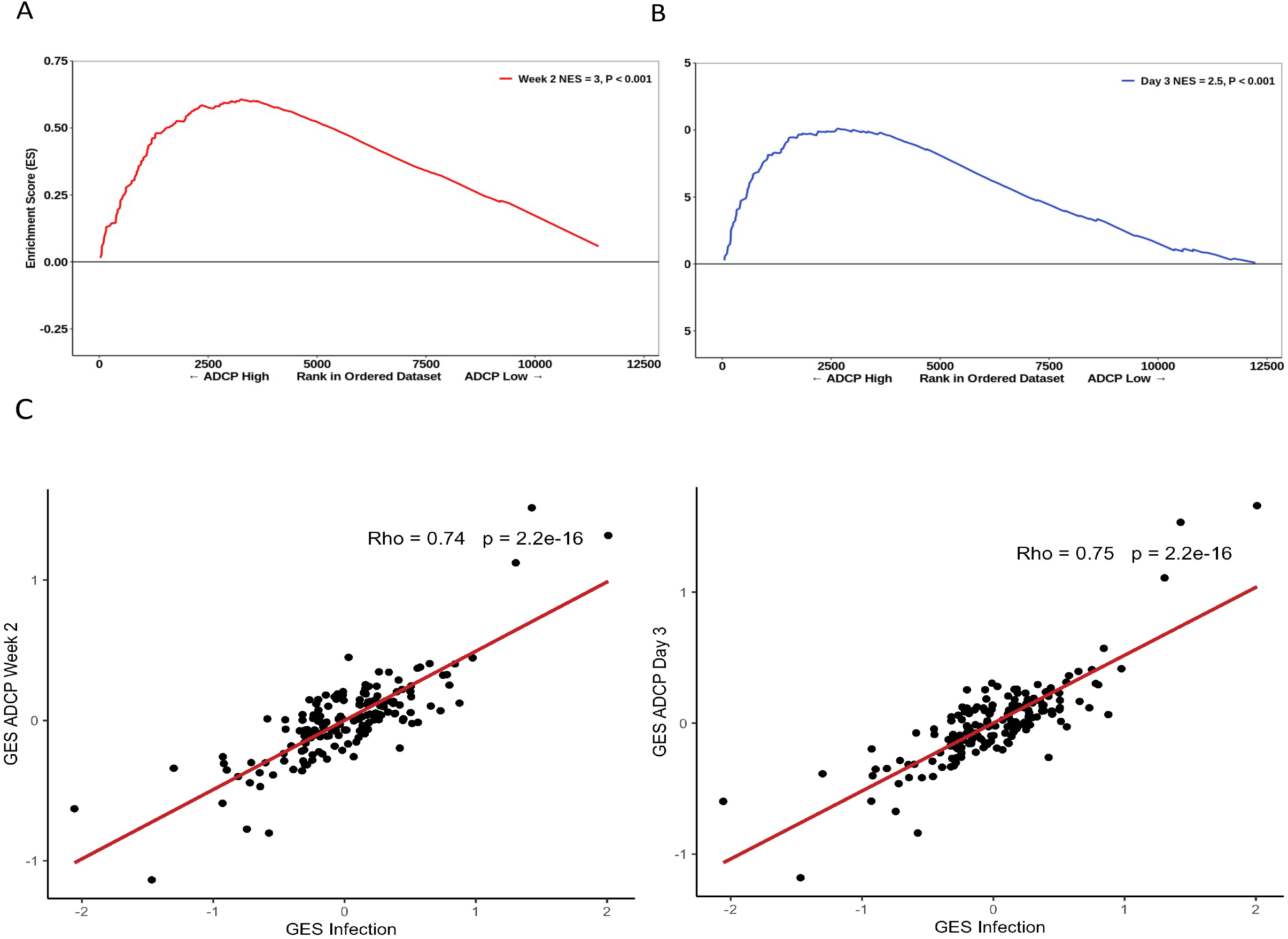
Strong relationship between functional ADCP responses in a human vaccine trial and the protective RV144 signature. The geneset that associated with protection in an efficacy study was also enriched with higher magnitude of ADCP measured two weeks after vaccination in an immunogenicity trial that employed the RV144 vaccine regimen. NES from RNA-seq data at timepoints (A) 2 weeks (118 enriched genes) (N = 24) and (B) 3 days (93 enriched genes) (N = 21) post the RV144 vaccine regimen in the RV306 trial are indicated. (C) GES computed from the enriched genes associating with ADCP correlated strongly with the protective GES in the RV144 study (N =170).

### Pathways are shared between ADCP and vaccine protection phenotypes

These findings demonstrated a strong link of the geneset with both vaccine protection and ADCP in NHP and human studies. We sought to broaden our understanding of the relationship between the different enriched genes in the geneset and establish some of the top pathways with gene membership from the different studies. Genes that were significantly enriched with either the ADCP or infection phenotypes from the 09-11, 13-19, RV144, and RV306 studies (Supplementary table 1) were classified into statistically significant terms from different biological pathways and represented as networks. The top significant pathways were leukocyte activation involved in immune responses, lysosome function and signaling by interleukins (Fig. 4A-B). Gene ontology of the 63 genes in the RV144 signature revealed that the top non-redundant enriched clusters with gene membership were myeloid leukocyte activation, lysosome and cellular response to oxidative stress genes (Fig. 4C).

**Fig. 4.**
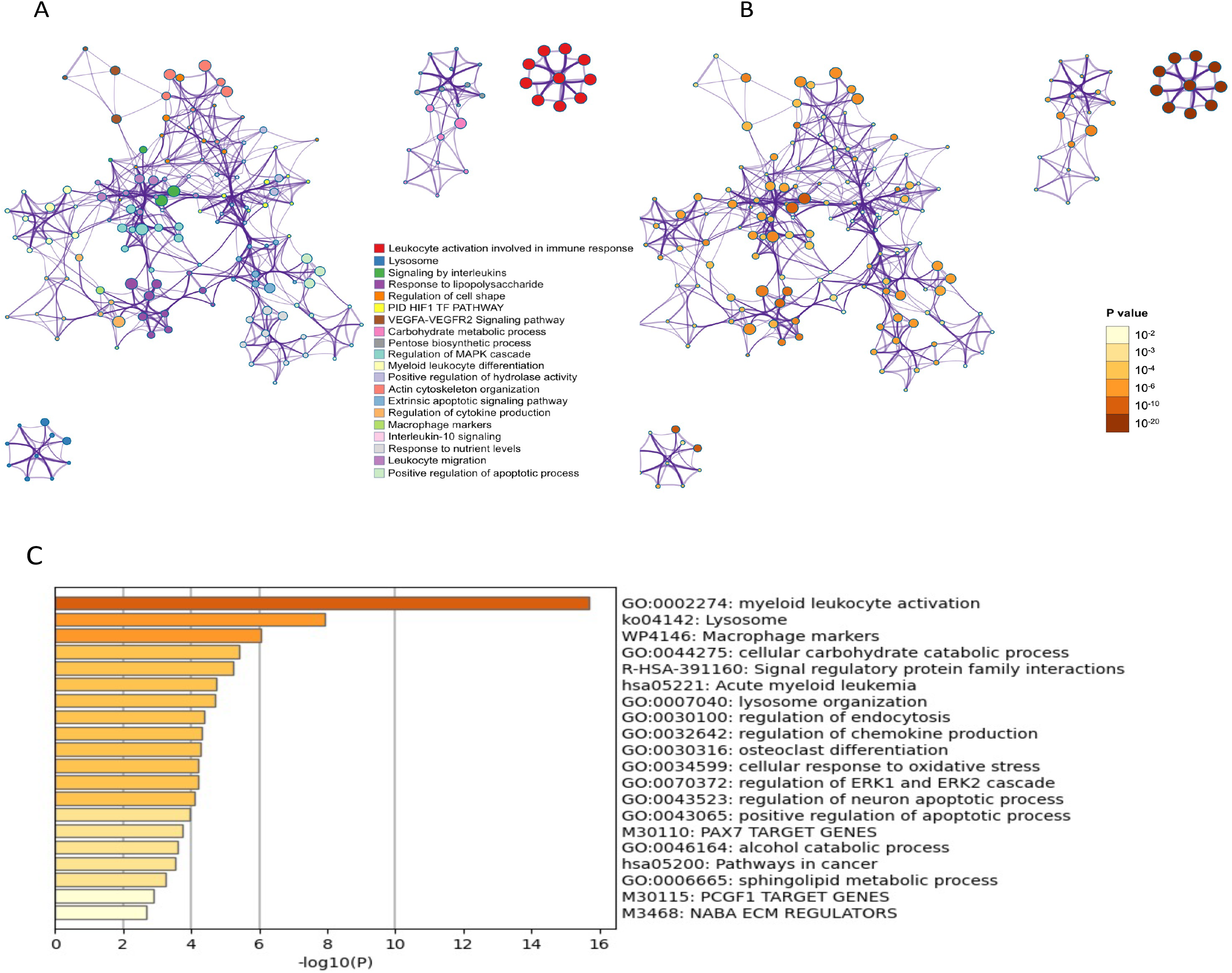
Pathway analyses of the enriched genes in the different vaccine studies. A meta-analysis of pathways including enriched genes with reduced infection or higher ADCP was performed. (A) The top related pathways and their (B) corresponding significance are indicated for the RV144 infection (N = 172), 09-11 (infection & ADCP) (N = 10), two arms of 13-19 (infection & ADCP) (N = 23), and RV306 ADCP (2 timepoints) (N = 45) enriched genes. Each term is represented by a circle node, where its size is proportional to the number of input genes that fall into that term, and its color represents its cluster identity. Terms with a similarity score > 0.3 are linked by an edge (the thickness of the edge represents the similarity score). The network is visualized with Cytoscape (v3.1.2) with “force-directed” layout and with edge bundled for clarity. (C) Clustering of the 63 enriched genes that associated with reduced infection in the RV144 study.

### Cellular origin of the protective genes by single cell transcriptomics

To dissect the cellular origin of these genes we performed simultaneous detection of mRNA and cell surface expression from single cells using the cellular indexing of transcriptomes and epitopes by sequencing (CITE-seq) technology in a subset of the vaccinated RV306 participants (Fig. 5A). This technology allows simultaneous detection of cell surface markers and gene expression from the same single cells. Our analysis revealed that a majority of the genes in the RV144 signature were expressed in cells of the myeloid lineage, with monocyte subsets having the highest average gene expression (Fig. 5B). A subset of genes were also significantly associated with decreased risk of acquisition in a univariate analysis (Odds ratio<1.0, P<0.05, q<0.1) (Fig. 5C). A stepwise logistic regression analysis identified specific genes (SEMA4A, SLC36A1, SERINC5, IL17RA, CTSD, CD68, GAA) to have independent associations with reduced risk of acquisition and were mainly expressed in the monocyte compartment (Fig. 5D). CD14 monocytes also had the greatest number of differentially expressed genes that were associated with ADCP (Fig. 5E).

**Fig. 5.**
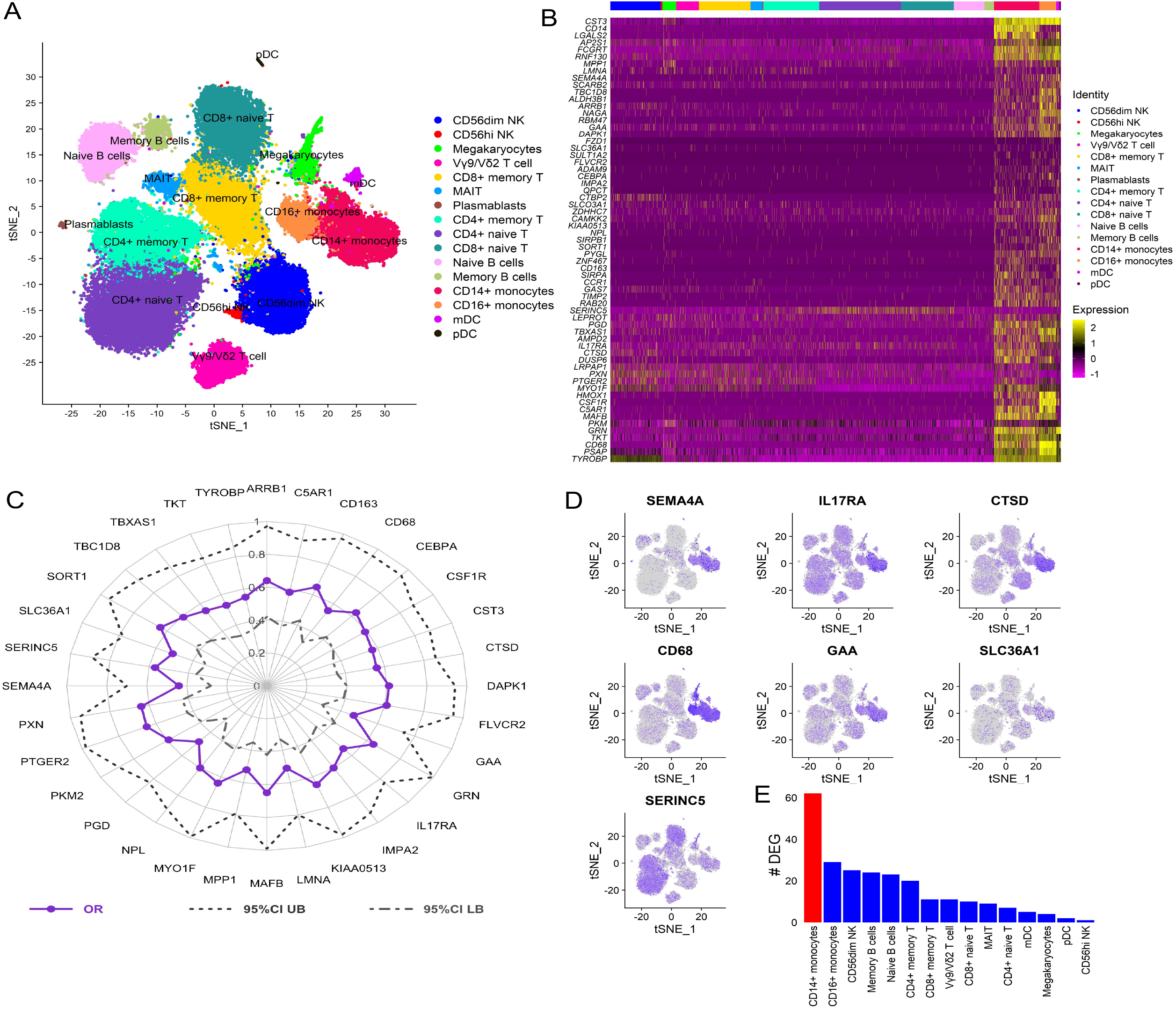
Cellular origin of the RV144 signature. Single cell CITE-seq in vaccinated participants (N = 12) that employed the RV144 vaccine regimen (day 3 after last vaccination) identified expression of the genes in the signature in cells from the myeloid lineage. (A) Clustering based on cell surface expression of CITE-seq data (B) Heat map of the mRNA expression of the 63 genes from the RV144 signature from single cells. Columns represent single cells from different protein cell subsets and rows the mRNA gene expression (C) Radar plot showing significant genes in the signature that associated with decreased risk of infection in RV144 (P<0.05, q<0.1) (N =170). (D) Feature plots of the expression of the most protective genes show that SEMA4A, IL17RA, CTSD, CD68 and GAA were mainly expressed in monocytes. (E) CD14+ monocytes had the highest number of differentially expressed genes (DEG) when comparing high versus low ADCP (2 weeks after vaccination) from single cell CITE-seq vaccinated participants that employed the RV144 vaccine regimen (day 3 after last vaccination).

## Discussion

Though an effective vaccine has been a challenge for the HIV field, we see a glimpse of optimism in partially protective NHP and human studies (Barouch et al., 2015; Barouch et al., 2013; Barouch et al., 2018; Rerks-Ngarm et al., 2009; Vaccari et al., 2018; Vaccari et al., 2016). These studies provide a unique opportunity to identify correlates of reduced risk that could help inform protective signals and enable design of enhanced vaccine strategies. Targeted and unbiased approaches have implicated non-neutralizing antibodies as the major correlate of reduced risk of HIV infection (Barouch et al., 2015; Barouch et al., 2013; Barouch et al., 2018; Haynes et al., 2012; Vaccari et al., 2018; Vaccari et al., 2016). We previously showed that a transcriptomic signature first identified in sorted B cells at timepoints prior to challenge was a correlate of protection in two NHP studies after administration of the Ad26/gp140 vaccine. This signature also associated with increased magnitude of ADCP in the vaccinated monkeys (Ehrenberg et al., 2019). We also identified this signature in bulk PBMCs from other studies that used the ALVAC/protein regimen, suggesting that this vaccine might be an indicator of effective vaccination. In this report we further investigated this gene signature to answer the following questions including: 1) can the gene signature’s association with protection be substantiated in additional human efficacy trials, 2) does it associate with ADCP in human trials, and 3) what is the cellular origin of the signature at the single cell level?

The gene signature previously associated with HIV vaccine protection in a number of studies with partial protection (Ehrenberg et al., 2019). We hypothesized that if this gene set was a true marker of HIV vaccine protection, it would not be enriched in a failed vaccine trial. HVTN 505 is a DNA based vaccine which despite not showing overall efficacy in a Phase 2b trial, demonstrated both cellular and antibody effector mediated protection in specific subgroups of individuals in follow-up studies (Fong et al., 2018; Janes et al., 2017; Neidich et al., 2019). We performed transcriptomics on sorted cell subsets from HVTN 505 vaccinated individuals and did not observe enrichment of the gene signature, further strengthening our notion that the geneset could be a proxy for vaccine protection.

Next, we developed a method to assess this gene signature compared to other correlates of risk in the human RV144 study. This method employs an analytical method using a GES which is computational score generated from the average expression of all genes enriched in the signature and associating with a phenotype. This method was tested across different NHP studies and RV144 and showed consistent association with reduced risk of infection based on the study specific GES. The composite GES computed from RV144 consisting of the standardized expression of 63 genes had the strongest association with decreased risk of infection and increased efficacy. The RV144 GES was also able to accurately predict infection status in the study. This study shows that the GES composite score provides a robust analytical measurement to explore the effect of genes as a continuous variable in immune-correlates analyses, and that it could be applied to other ongoing efficacy studies. Further, the analysis of RV144 breakthrough infections were consistent with GES being lower in the vaccinated infected participants compared to placebos and hence protective. These observations albeit only significant in the group infected with single founder viruses, strengthen the premise of the RV144 GES being a correlate of reduced risk of infection.

We previously showed in NHP challenge studies that the gene signature correlated with increased magnitude of functional antibody responses (Ehrenberg et al., 2019). Although this geneset associated with ADCP in in NHP, the same analyses were previously not possible in the human RV144 study since this immune response was not reported (Haynes et al., 2012). ADCP has since been implicated with vaccine protection in a number of NHP challenge studies (Ackerman et al., 2018; Barouch et al., 2015; Barouch et al., 2013; Barouch et al., 2018; Bradley et al., 2017; Neidich et al., 2019). It is reported that ADCP could be involved in most studies that previously showed antibody-dependent correlates of protection against viruses (Tay, Wiehe, & Pollara, 2019).To investigate the effect of the gene signature on the magnitude of ADCP, we performed transcriptomics in samples from a human trial (RV306) that employed the same RV144 regimen. At both day 3 and two weeks after the 4^th^ vaccine corresponding to the last RV144 vaccine dose, this signature correlated with increased magnitude of ADCP responses. A strong correlation was also observed between GES from the ADCP enriched genes and the vaccine protection genes in RV144. These findings provide evidence supporting the antibody-mediated effector function as the mechanistic basis of this signature.

The enriched genes from ADCP and infection risk in multiple studies of both NHP and human were involved in overlapping functions related to leukocyte activation, lysosomal degradation and immune stimulation by cytokines. The 63 genes from the RV144 signature were mostly clustered in the myeloid leukocyte activation pathway, alluding to the cellular origin of this signature. The specific genes in the gene set that associated with the greatest odds of reduced risk of infection including SEMA4A, CTSD, CD68 and GAA were all members of this pathway, but not TNFSF13 (APRIL) which was the most protective gene in the NHP studies. Although the geneset of interest was first seen in sorted B cells from vaccinated NHP, it was subsequently identified in transcriptomic data from PBMCs in the RV144 study (Ehrenberg et al., 2019). While samples were exhausted from the RV144 primary dataset, the RV306 clinical trial that employed the same ALVAC-protein vaccine regimen gave us a unique chance to explore the cellular origin of the RV144 signature using single cell transcriptomics. Single cell surface expression data revealed that the majority of genes were expressed in monocytes, which was not surprising given the fact that this geneset was originally defined as genes downregulated in B cells compared to monocytes after influenza vaccination (Nakaya et al., 2011). While our initial study found this signature in sorted B cells from the NHP challenge studies, single cell data provides further insight that monocytes could be the cellular origin of these genes in the RV144 study. Although monocytes were classified as mononuclear phagocytes almost fifty years ago, assays designed to specifically measure monocyte ADCP were not widely used in the context of vaccination until a few years ago (van Furth et al., 1972). Although monocytes have recently been implicated in vaccine-induced protection in preclinical vaccine trials of SIV challenge, our findings in human trials at the single cell level provides greater impetus to explore the role of other non-lymphoid cell populations on HIV-1 vaccine efficacy. (Gorini et al., 2020; Vaccari et al., 2018). Though we think that monocytes are important in the vaccine responses observed in RV144, it would be remiss not to mention that the effect of granulocytes including neutrophils in response to vaccination is missed when transcriptomics is performed in PBMC compared to blood. Other than the phagocytic cell, both antibody and Fc receptor diversity can influence ADCP mediated immune responses to viral pathogens and are also elements that warrant further study and may potentially be manipulated to improve vaccine efficacy (Chung & Alter, 2017; Geraghty, Thorball, Fellay, & Thomas, 2019; Tay, Wiehe, et al., 2019).

Our data demonstrate the potential to discover novel protective correlates using an approach that mines transcriptomic data in multiple preclinical and clinical trials. Unbiased transcriptome-wide analyses are able to identify biological perturbations that associate with vaccine protection even when differences are small, but credibility can only be strengthened by replicating findings across multiple studies. Gene signatures that associate not only with vaccine protection, but specific immune responses can be a prospective tool to evaluate vaccine effectiveness even prior to challenge or infection. Developing analytical tools that can interface with phenotypes such as vaccine protection across human and preclinical studies can allow for more systematic meta-analyses of data emerging from the ongoing HIV vaccine clinical trials, as well as the recently halted HVTN 702 trial (NIH, 2020). We propose that assessment of such gene signatures with immune responses in human immunogenicity trials could provide orthogonal insight for down-selection of vaccine candidates. Identifying overlapping correlates of protection in these studies could be pivotal to making discoveries that may allow for licensure and subsequent bridging studies of an effective HIV vaccine.

## Methods

### Study design

The aim of the study was post-hoc analyses of a protective gene expression signature identified previously in five SIV/HIV vaccine studies with efficacy and immune response data (Ehrenberg et al., 2019). To enable interpretation of this gene signature, bulk RNA-seq, scRNA-seq and functional data was generated in clinical samples from the RV306 and HVTN 505 human trials. The RV306 vaccine trial was conducted in Thailand and all participants received the primary RV144 ALVAC/gp120 vaccine series, with additional late boosts assigned to specific groups (Pitisuttithum et al., 2020). Bulk RNA-seq was performed in 24 participants two weeks after the RV144 vaccine regimen (week 26). Additionally, RNA-seq was also performed 3 days after the same primary endpoint. The HVTN 505 trial used a DNA/rAd5 vaccine regimen to test safety and efficacy in a US population (Hammer et al., 2013). PBMC collected one month after the final immunization (month 7) was available from 47 vaccinees in the HVTN 505 study for RNA-seq (Hammer et al., 2013). The infection status of the vaccinees (22 cases and 25 controls) was categorized based on infection status between months 7-24. Microarray transcriptome data from PBMCs and immune response data for 170 vaccinated individuals from the RV144 study at timepoint two weeks post last vaccination was used for correlates analyses (Fourati et al., 2019; Haynes et al., 2012). All studies were approved by the participating local and international institution review boards. Informed consent was obtained from all participants in the different trials included in this study (Hammer et al., 2013; Pitisuttithum et al., 2020).

### Bulk transcriptomics

RNA was extracted from sorted B cells (Aqua live/dead^−^CD20^+^CD3^−^) and monocytes (Aqua live/dead^−^CD20^−^CD3^−^CD56^−^HLA-DR^+^CD14^+^ from PBMC of HVTN 505 vaccinees using RNAzolRT (MRC Inc.) as per recommendations from the manufacturer. For the preparation of mRNA libraries, polyadenylated transcripts were purified on oligo-dT magnetic beads, fragmented, reverse transcribed using random hexamers and incorporated into barcoded cDNA libraries based on the Illumina TruSeq platform. Next, libraries were validated by electrophoresis, quantified, pooled and clustered on Illumina TruSeq v2 flow cells. Clustered flow cells were sequenced on an Illumina HiSeq (2000/4000) using 2 × 75 base paired-end runs. Total RNA from RV306 participants was extracted from whole blood collected in PAXgene Blood RNA tubes, using the PAXgene Blood RNA kits (both Qiagen; Germantown, MD) and subsequently subjected to the GlobinClear kit (ThermoFisher Scientific; Waltham, MA) per manufacturer’s suggestions. NGS RNA-seq was performed using the SMART-Seq technology (Picelli et al., 2014; Ramskold et al., 2012). Briefly, cDNA was generated from 10 ng of RNA using the SMART-Seq v4 UltraLow Input RNA Prep kit (Takara Bio Inc) per manufacturer’s suggestions, with control RNA spiked-in (ThermoFisher Scientific). Sequencing libraries were generated using the Nextera XT DNA Sample Prep kit (Illumina, San Diego, CA). Concentration of each sample in the pooled libraries were determined using the paired-end 300-cycle MiSeq Reagent Nano Kit v2 (2 × 150 bp) on a MiSeq instrument (both Illumina). NGS was performed on a final adjusted library pool using the paired-end 300-cycle NovaSeq 6000 S2 XP Reagent Kit (2 × 150 bp) on a NovaSeq instrument (both Illumina) per the manufacturer’s instructions. Fastp v0.19.7 and Trimmomatic v0.33 with default parameters was used to trim low-quality bases from both ends of each read. Trimmed reads were aligned to the human genome (GRCh38 build 88-92) using HISAT2 v2.1.0 or the STAR aligner (v2.4.2a) and HTSeq (v0.6.1-0.9.1) was used for counting. Trimmed mean of M-values normalization method, as implemented in the R package edgeR, was used for normalization.

### Single cell transcriptomics

Simultaneous evaluation of mRNA and cell surface expression from single cells was performed using feature barcoding technology from 10x Genomics, based on the CITE-seq technology (Stoeckius et al., 2017). Cell hashing (HTO) was used in conjunction with the 10x Genomics 5’V(D)J Feature Barcoding kit to generate single cell mRNA (GEX) and antibody-derived tag (ADT) libraries (Stoeckius et al., 2017; Stoeckius et al., 2018). Briefly, PBMCs from 12 samples were hashed using TotalSeq-C anti-human Hashtag antibodies and combined into two batches. In each batch, surface proteins were stained with a cocktail of 53 TotalSeq-C antibodies (Biolegend). Antibody concentrations were either predetermined by titration (Kotliarov et al., 2020) or used at a default concentration. 50,000 cells from each batch were loaded onto each of 4 wells of a Chromium Chip, and GEX and ADT (HTO and Feature Barcode (FB)) libraries were constructed following the manufacturer’s protocol. Libraries were pooled and quantitated using a MiSeq Nano v2 reagent cartridge. Final libraries were sequenced on the NovaSeq 6000, S4 reagent cartridge (2×100 bp) (Illumina).

### CITE-seq data analyses

FASTQ files were demultiplexed with bcl2fastq v2.20 (Illumina). Alignment and counting were performed using Cell Ranger v3.1.0 (10x Genomics) and the human reference files provided by 10x Genomics (human genome GRCh38 and Ensembl annotation v93). The average number of genes per cell was 1453 and the average number of unique molecular identifiers (UMI) was 4248. The mean read depth per cell was approximately 65,000–84,000. The minimum fraction of reads mapped to the genome was 88% and sequencing saturation was above 85% for all lanes with an average of 88%. The computational analysis of ADT data was performed using the Seurat v3.1 package (Stuart et al., 2019). HTO expression matrices were CLR (Centered Log-Ratio) normalized and demultiplexed using MULTIseqDemux. The FB matrices from the Seurat objects were split into cell-positive and negative droplet matrices using the HTO demultiplexing results, and were used for denoising by DSB (Denoised and Scaled by Background) normalization (Kotliarov et al., 2020)(https://cran.r-project.org/web/packages/dsb/index.html). Only cells with <10% mitochondrial genes were retained, and cells were assigned to specific donors using the HTO demultiplexing results. A total of 53,777 single cells remained after the quality control process. The gene expression matrices for all samples were normalized and integrated into a single object in Seurat (Stuart et al., 2019). Based on the workflow described in Kotliarov et al., a distance matrix was generated from cell surface protein features (Kotliarov et al., 2020). This matrix was used for shared-nearest-neighbor finding and clustering at resolution=0.5. Neighbor finding and clustering were performed on the integrated gene expression data at a resolution=0.75 and dimensions=1:30. A tSNE (t-distributed stochastic neighbour embedding) was generated from the protein data PCA. Seurat was used to generate a heatmap, dotplot and featureplots. Differential gene expression testing was performed within each cluster between the high and low ADCP groups using Seurat’s FindMarkers function. ADCP DEG were filtered to genes with >10% expression in either group, a log fold change value > 0.25, and a Bonferroni p value < 0.05.

### ADCP assay

The antibody effector function ADCP was measured as previously described (Ackerman et al., 2011; Tay, Kunz, et al., 2019; Tay et al., 2016). Briefly, A244 gp120 Env-coated fluorescent beads were incubated at 37°C for 2 hours with diluted plasma (1:50) collected at week 26, two weeks after administration of the RV144 vaccination series. Anti-CD4 monoclonal antibody-treated THP-1 cells (human monocytic cell line; ATCC TIB-201) (treated for 15 minutes at 4°C) were added to immune complexes and spinoculated for 1 hour at 4°C to allow phagocytosis to occur. Supernatant was removed, cells were washed and fixed in paraformaldehyde. Phagocytosis was measured by flow cytometry and a phagocytosis score was calculated as follows: phagocytosis score = (% pos * MFI of Sample) / (% pos * MFI of no-antibody PBS control). The CD4 binding-site broadly neutralizing antibody (bnAb), CH31, was used as a positive control, and the influenza receptor binding site-specific bnAb, CH65, was used as a negative control. Results are representative of two independent experiments.

### Pathway analyses

Association of the protective gene signature with infection (HVTN 505) or magnitude of median ADCP (RV306) responses were analyzed using the Gene Set Enrichment Analysis (GSEA) method as described previously (Ehrenberg et al., 2019; Subramanian et al., 2005). GSEA was performed on vaccinated HVTN 505 participants at the visit 7 timepoint, one month after the last immunization. RNA-seq was performed on samples prior to infection, but participants were categorized based on their infection status. GSEA was performed on 45 RV306 RNA-seq samples that also had ADCP scores obtained at the week 26 (week 2 after the 4^th^ vaccination) time-point. Participants were categorized into high and low ADCP groups based on the median values of ADCP measured in a total 79 vaccinated participants. The RNA-seq gene expression values at the day 3 and week 2 time-points were then analyzed for gene enrichment using a gene set of 200 genes, obtained from the Broad Institute (GSE29618_BCELL_VS_MONOCYTE_DAY7_FLU_VACCINE_DN), between the two groups of samples based on the classification, in each time-point. The gene signature of interest was considered significantly enriched using a threshold of NES≥1.4 and P<0.001 as described previously (Ehrenberg et al., 2019). Gene ontology and network analyses of genes were performed using Metascape with default parameters (Zhou et al., 2019).

### Correlates of protection

Composite gene expression score (GES) was computed as the average of standardized expression of normalized enriched genes in the gene signature in different vaccine studies. The samples in each vaccine study were grouped into outcomes after challenge or infection status after immunization (Barouch et al., 2015; Barouch et al., 2018; Rerks-Ngarm et al., 2009). Logistic regression was used for evaluating the association between GES and HIV-1 infection in the RV144 study. The fitting methods accommodate the 2-phase sampling design via maximum likelihood estimation (Breslow & Holubkov, 1997). Cumulative HIV-1 incidence curves were plotted for the three subgroups of vaccine recipients defined by tertiles into the lower, middle, and upper third of the GES (Low, Medium, High subgroups), as well as for the entire placebo group HIV negative at week 24 (n=6267 subjects) for reference. These curves were estimated using the Kaplan-Meier method with inverse probability weighting that accounted for the sampling design. Next, vaccine efficacy (VE) for the GES subgroups versus the entire placebo group was estimated as one minus the odds of infection in vaccine recipients with Low/Medium/High response divided by the odds of infection in the entire placebo group HIV-1 negative at week 24 of enrollment in the study. The RV144 prediction analysis was implemented by logistic regression. The dataset was randomly split into training and testing sets in a 7:3 ratio, while retaining class distributions within the groups. The training dataset consisted of 119 individuals while the test dataset consisted of 51 individuals. A logistic regression of GES was fit on to the training dataset (Prentice et al., 2015). The model’s discriminative ability was evaluated by generating a receiver operator characteristic curve (ROC) and the corresponding AUC on the test dataset. The prediction accuracy of the model was also assessed on the test dataset. The probability that gives minimum mis-classification error was chosen as the cut-off. This process was repeated 1000 times and the distribution of the resulting AUC and accuracy were demonstrated by a histogram with a density curve.

Among 121 RV144 participants who became infected during the trial and had their HIV-1 genome sequenced at diagnosis, 43 had GES measurements computed from microarray data (Fourati et al., 2019; Rolland et al., 2012). Vaccine and placebo groups were compared overall and after stratifying infections with single HIV-1 founders.

### Other statistical analyses

Logistic regression that accounted for the sampling design was used for the univariate analyses of the 63 enriched genes. A radar plot of the significant genes was generated to illustrate odds ratios (OR) and 95% confidence intervals. All ORs were reported per 1-SD increase. Significant genes resulted from univariate logistic regressions of the 63 enriched genes were further analyzed with a multivariate stepwise logistic regression to identify genes that independently associated with HIV protection. Akaike information criterion (AIC) was used to identify the optimal set of genes. The expressed enriched genes associated with higher magnitude of ADCP in RV306 at day 3 and 2 weeks post the RV144 vaccine regimen were used to compute the ADCP GES in RV144. Spearman correlation was calculated between the ADCP GES from the two timepoints and the infection GES respectively.

All descriptive and inferential statistical analyses were performed using GraphPad Prism 8 (GraphPad Software) and R 3.6.1 (or later) software packages. Comparison of groups was performed using Mann-Whitney tests or t-tests when assumptions were met. All logistic regression models were adjusted for gender and baseline risk behavior and one significant principal component axis (Haynes et al., 2012; Prentice et al., 2015). A two-sided P value of less than 0.05 was considered significant. The Benjamini and Hochberg method was used to calculate false discovery rate (FDR)-adjusted p values for multiple testing corrections.

## Acknowledgements

We would like to thank the volunteers and staff of the RV306, RV144 and HVTN 505 clinical trials. We also acknowledge DeAnna Tenney and Derrick Goodman, Duke University for expert technical assistance. The views expressed are those of the authors and should not be construed to represent the positions of the U.S. Army or the U.S. Department of Defense (DOD). This work was supported by a cooperative agreement (W81XWH-07-2-0067) between the Henry M. Jackson Foundation for the Advancement of Military Medicine, Inc., and the DOD. This research was funded in part by the U.S. National Institute of Allergy and Infectious Disease.

## Supplementary Material

**Supplementary Fig. 1.**
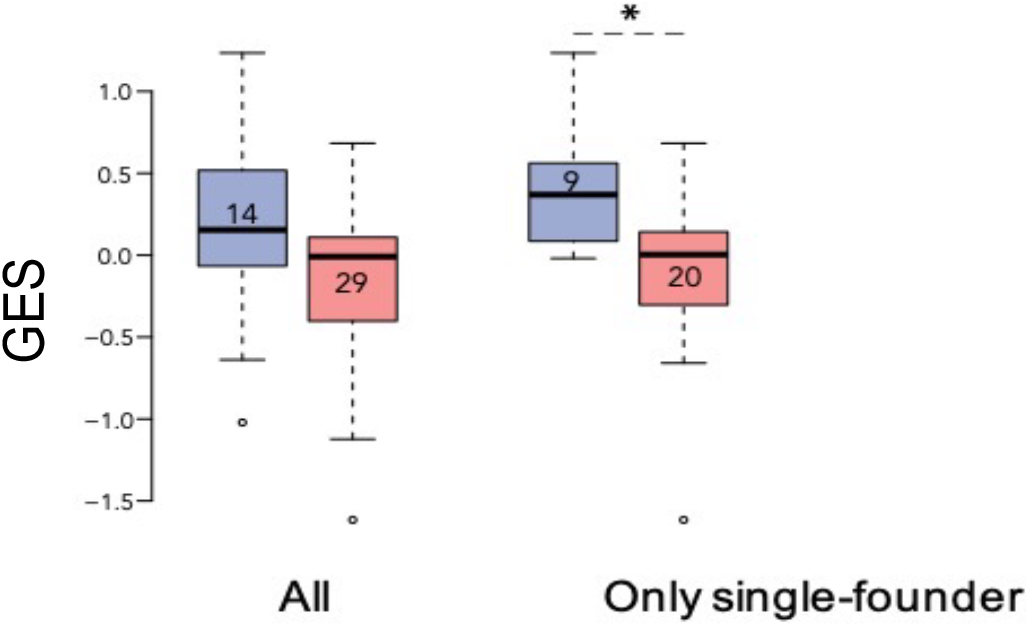
Association of the GES with HIV-1 breakthrough infections in a human vaccine trial. GES was significantly higher in the placebo (blue) compared to vaccine recipients (red) among the RV144 participants with single-founder breakthrough infection (P<0.05). Numbers indicate the number of participants plotted. Asterisks indicate significant pairwise differences by Mann-Whitney test.

**Supplementary table 1.**Number of enriched genes from the geneset in different comparisons from multiple studies.

